# Cryptic female choice in response to male pheromones in *Drosophila melanogaster*

**DOI:** 10.1101/2023.12.20.572608

**Authors:** Nicolas Doubovetzky, Philip Kohlmeier, Sanne Bal, Jean-Christophe Billeter

**Affiliations:** Groningen Institute for Evolutionary Life Sciences, University of Groningen, 9474AG Groningen, The Netherlands; University of Memphis, Department of Biological Sciences, Memphis, TN 38152-3530, USA

**Keywords:** Cryptic female choice, post-mating sexual selection, sperm ejection, sperm storage, pheromones, olfaction, *Drosophila melanogaster*

## Abstract

Females control the paternity of their offspring by selectively mating with males they perceive to be of high quality. In species where females mate with multiple males in succession, females may bias offspring paternity by favoring the sperm of one male over another, a process known as cryptic female choice (CFC). While evidence of CFC exists in multiple taxa, the mechanisms underlying this process have remained difficult to unravel. Understanding CFC requires demonstration of a female-driven post mating bias in sperm use and paternity, and a causal link between this bias and male cues. Here, we show that in the vinegar fly *Drosophila melanogaster,* mated females eject the ejaculate of their first mate faster when exposed to the pheromones of an attractive male than in the presence of an unattractive one. Using transgenic males expressing fluorescent sperm, we show that exposure to attractive males between mating causes twice-mated females to bias sperm storage towards the second male, affecting paternity. Using pheromonal bioassays in combination with genetic manipulation of sensory systems, we show that females modulate ejaculate ejection latency in response to male pheromones heptanal and 11-cis-Vaccenyl acetate (cVA) sensed via olfactory receptor neurons OR35a, Or22a, Or65a and OR67d, demonstrating that polyandrous females use male pheromonal cues to modulate ejaculate ejection timing. We provide the first demonstration to our knowledge of a CFC mechanism allowing a female to increase or decrease the share of paternity of her first mate depending on the sensing of the quality of potential mates in her environment. These findings showcase that paternity can be influenced by events that go beyond copulation and highlights the importance of post-copulatory sexual selection.

**One Sentence Summary:** We show that females bias sperm use and paternity towards specific males in response to male pheromones.

## Introduction

Individuals of sexually reproducing species evolved multiple mechanisms to identify and select high quality mating partners. These traits include pre-copulatory mechanisms such as selecting highly fecund mates, or those with genetic quality that confer survivability and fecundity to their offspring. In species where females mate with multiple males in one reproductive episode, sperm from different males interact in the female reproductive tract. This creates an opportunity for competition between sperms, often leading the last male to mate to father most of the offspring - a phenomenon known as last male sperm precedence. Although often overlooked, polyandry also allows for females to apply post-copulatory mechanisms that bias sperm use towards one male or the other, affecting the share of offspring of each male – a poorly understood phenomenon called cryptic-female choice (CFC).

One CFC mechanism is ejaculate ejection, via which females expel sperm from a recent mating which can bias that male’s paternity share (Eberhard et al., 1996; Eberhard et al., 2009; Firman et al., 2017). In the feral fowl, females have a higher chance of ejecting the ejaculate of a low-ranking male than a high-ranking one, which favors paternity by higher ranking males (Pizzari & Birkhead, 2000). In the Ulidiid flies *Euxesta bilimeki*, females partially or totally expel sperm transferred during mating depending on their mate size (Rodriguez-Enriquez et al., 2013). Understanding the mechanisms that underlie CFC has however remained elusive because this process happens hidden in the female reproductive tract and requires demonstration of a female-driven post mating bias in sperm use and paternity, and a causal link between this bias and male cues (Firman et al., 2017).

*Drosophila melanogaster* is an ideal model to investigate CFC mechanisms because it is a polyandrous species (Billeter & Wolfner, 2018; Firman et al., 2017) with extant genetic tools labelling spermatozoa of different males with different fluorescent markers allowing for direct observation in the female reproductive tract (Laturney et al., 2018; Lee et al., 2015; Manier et al., 2010). Moreover, the cues based on which females choose which male to mate with, a pre-copulatory choice, are also partially identified, including the male pheromones cis-Vaccenyl Acetate (cVA), Palmitoleic acid, and 7-Tricosene (Billeter et al., 2009; Grillet et al., 2006; Kohlmeier et al., 2021; Kurtovic et al., 2007). *D. melanogaster* females also naturally eject male ejaculates. Indeed, directly after mating females store a portion of the sperm from the male ejaculate in sperm storage organs located in their reproductive tract, a process that lasts a few hours, and then eject the unstored ejaculate from their reproductive tract (Laturney & Billeter, 2016; Lee et al., 2015; Manier et al., 2010). After this first ejaculate ejection, females can remate with a second male, which occurs in both lab and natural settings (Billeter & Wolfner, 2018; Laturney & Billeter, 2016). With the arrival of the new ejaculate, second male sperm displaces a portion of the first male sperm from the female sperm storage organs, and these displaced sperms are then ejected by the female (Manier et al., 2010). The timing of this second ejaculate ejection correlates with paternity in a way where the longer females take to eject, the more displacement can occur, and the more eggs are sired by the second male (Lüpold et al., 2013). Although the mechanism remains unknown, it is presumed that in this case, females have more time to store sperm from the second male and for it to displace resident sperms from the first male.

Given that ejaculate ejection latency is associated with female sperm storage, we hypothesized that it could be a CFC mechanism reducing use of sperm from a recent mate in the presence of attractive males in order to favor the sperm of these potential future mates. In this study, we reveal an effect of male quality on the timing of ejaculate ejection and offspring production in *D. melanogaster*. We show that females eject the ejaculate from their first mate earlier in the presence of an attractive male. When females remate, the advanced ejection correlates with a bias in paternity and sperm stored by the female, biasing paternity by the second male. We also identify that the male pheromones cVA and heptanal modulate these effects through detection by specific female odorant receptors. We thus propose a mechanism of cryptic female choice that bias paternity in response to male cues.

## Results

### Mated females modulate ejaculation latency and bias paternity depending on the quality of potential mates in their environment

Cryptic female choice (CFC) is a post-copulatory mechanism that biases sperm use and impacts paternity ratios. We hypothesized that female modulation of ejaculate ejection latency is a CFC mechanism, allowing females to bias the paternity share of a given male depending on the female’s mating opportunities. If true, we predicted that recently mated females should shorten the ejection latency of their first mate ejaculate when sensing a preferred mate in their environment, resulting in a decrease in first mate share of paternity in favor of their next mate.

To identify preferred males, we assessed *Canton-S* (*CS*) female mate choice between males from two strains*, CS* and *Oregon-R* (*OR*). In a competitive mate choice experiment, we found that 82.7% of virgin females mated with *CS* over *OR* males (n=53; Fisher’s exact test; P<0.0001, Supp fig 1A). Next, to link preference with ejection latency, we tested if mate preference also impacted the ejaculate ejection latency given different postcopulatory conditions. We exposed single *CS* females recently mated to a *CS* male to either a *CS*, *OR*, or no male and measured ejaculate ejection latency. We found that females that were exposed to *CS* males ejected about 60 minutes earlier compared to females exposed to *OR* males or females kept in isolation (Fig 1B). Recently mated females thus advance timing of ejaculate ejection from their first mate only when exposed to preferred males.

**Figure 1.**
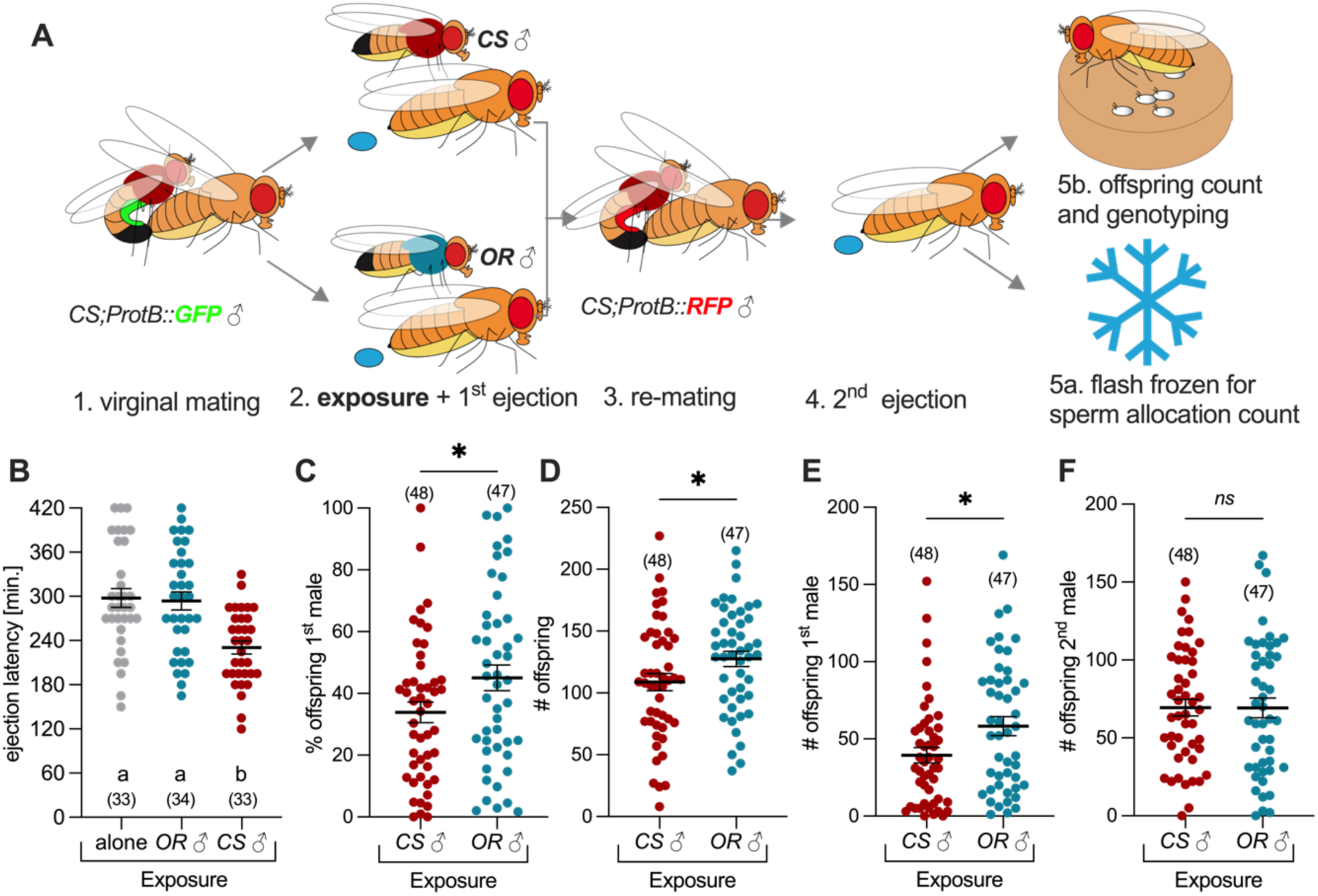
Twice-mated females modulate ejaculate ejection latency and bias paternity depending on the quality of males they are exposed immediately after their first mating. (A) Schematic of the experiment. (B) Ejaculate ejection latency of mated *Canton-S* (*CS*) females isolated, in the presence of an *Oregon-R* (*OR*) male or a *CS* male. (B-F) Offspring production of *CS* females twice mated with CS males as in (A) but exposed to either one *CS* or *OR* male in-between the two mating: (C) Percentage (%) of offspring from the first male, (D) Total number (#) of offspring, (E) Number of offspring from first male, and (F) Number of offspring from second male (mean ± S.E.M). Numbers between brackets indicate sample size. Mean ± S.E.M are indicated in the graphs. Scatter-plot labelled with the same letter are not statistically different. *n.s*.: non-significant; *:p<0.05; **:p<0.01; ***:p<0.001.For full statistical details see Supplementary Table S1.

To test whether modulation of ejaculate ejection latency for the first mate impacts paternity share, females were first mated with a *CS* male with Green Fluorescent Protein (GFP)-labelled sperm, then exposed to either *CS* or *OR* males until the first ejaculate ejection (before they could remate) and then immediately transferred in an arena with a *CS* male with Red Fluorescent Protein (RFP) labelled sperm for remating (Fig1.A). This produced double-mated females that either advanced or did not advance ejaculate ejection time after their first mating but otherwise mated with similar males, removing confounding factors such as differences in male sperm competitive abilities. Females exposed to *CS* or *OR* males between first and second mating ejected ejaculate from their second mates at similar times, indicating that all variation comes from the period during which once mated females were exposed to *CS* or *OR* males (Supp Fig 1F). These twice-mated females were then either isolated and the paternity of the resulting offspring determined to belong to the first or the second *CS* male based on GFP/RFP expression, or flash frozen to determine the number of sperms stored from GFP/RFP males (Fig 1A). We found that twice mated females exposed to *CS* males between first and second mating produced a lower proportion of offspring from their first mate than females exposed to *OR* males (Fig 1C). These data show that exposure to a preferred male after the first mating resulted in lower first mate paternity share. Females exposed to high quality *CS* males between first and second mating had an overall lower fecundity (Fig 1D) and, critically, less offspring from the first male compared to females exposed to *OR* males (Fig 1E), while the number of offspring from the second male remained similar in the two exposures (Fig 1F). These results show that *D. melanogaster* females bias paternity in favor of the second male when exposed to high-quality males in their post mating environment and, alternatively biasing paternity in favor of the first male when sensing low-quality mates. We thus show that *D. melanogaster* females can exert CFC in response to the presence of preferred males.

### Females bias sperm storage in the seminal receptacle depending on male quality

Females store sperm in two different specialized structures in their reproductive tract. The seminal receptacle is the main organ storing the majority of sperm, while the spermathecae store less sperm but for longer term (Pitnick et al., 1999; Schnakenberg et al., 2011, 2012)(Fig 2A). To understand how females bias paternity, we examined the proportion of the first male sperm (labeled with GFP) and second male sperm (labeled with RFP) in storage in twice mated females exposed to *CS* or *OR* males between first and second mating (as in Figure 1A). For this, we counted sperms stored in the seminal receptacle and the spermathecae separately. Although the total number of sperm from both males in storage in the seminal receptacle was similar between treatments (Fig 2B), we observed a bias against the first male’s sperm when the female was exposed to a *CS* male, and in favor of the first male’s sperm in the presence of an *OR* male (Fig 2C). These biases result from both a lower number of first male’s sperm (Fig 2D) and a higher number of second male sperm (Fig 2E) stored in the seminal receptacle when exposed to a *CS* male, and the opposite when exposed to an *OR* male. When analyzing sperm storage in the spermathecae, we observed an equal amount of sperm between groups (Fig 2F), but no bias in the proportion of first versus second male sperm (Fig 2G). There were also no differences in the number of sperm from the first (Fig 2H) or second male (Fig 2I). Difference in sperm allocation in the presence of *OR* or *CS* males between two mating is thus in keeping with differences in paternity observed depending on male exposure in the seminal receptacle but not in the spermathecae (Fig. 1). We conclude that females exert CFC by biasing sperm storage specifically in the seminal receptacle.

**Figure 2.**
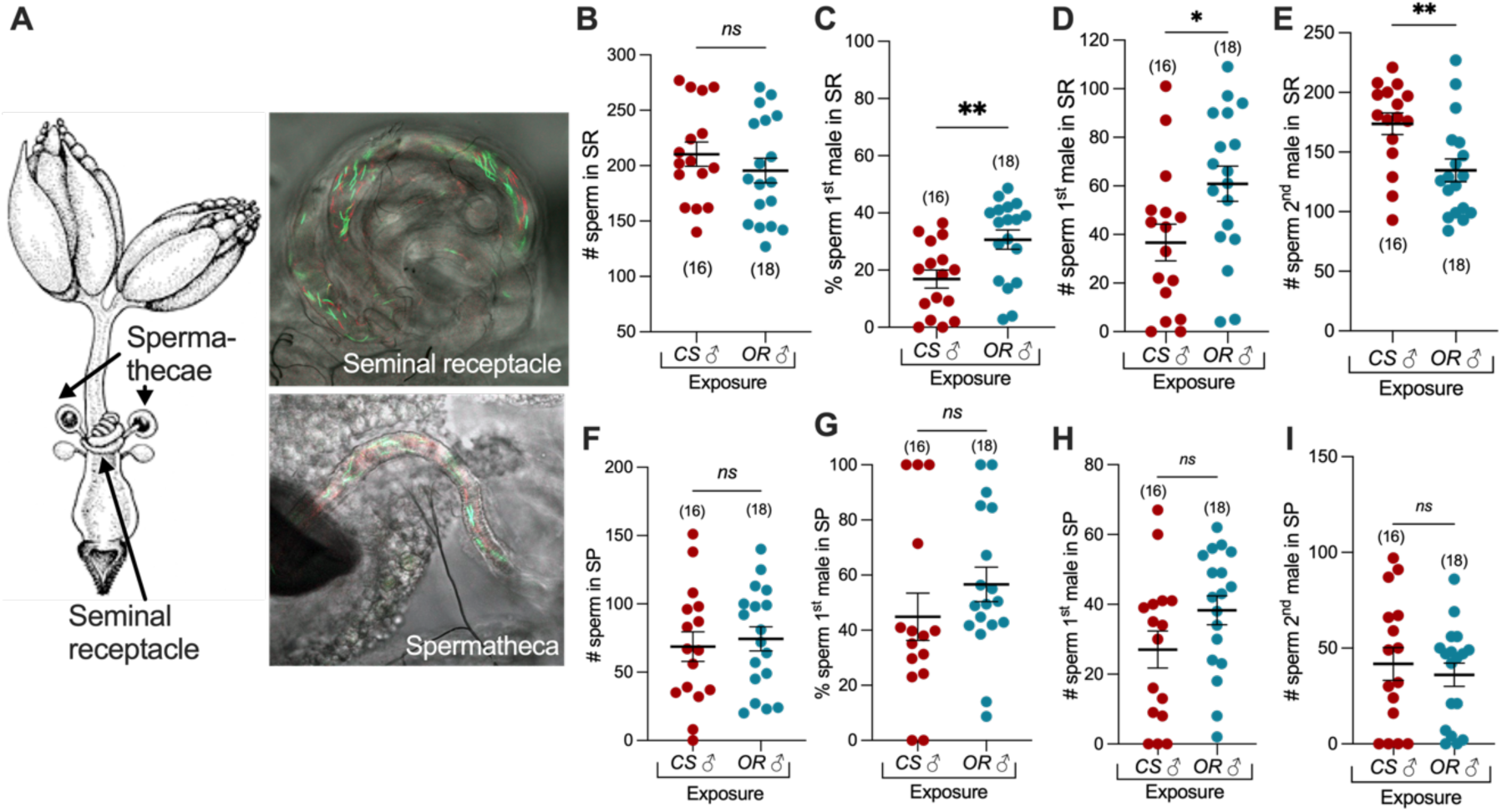
Sperm storage bias toward the first male when exposed to a low-quality potential mate between two mating. (A)Schematics of the female reproductive tract with confocal microscopy images detailed green and red sperms from different males in the seminal receptacle or the spermatheca. (B-I) number (#) or percentages (%) of sperms from first or second males in storage in the indicated sperm storage organ of a twice mated females exposed to either *CS* or *OR* males between the two mating (See Fig 1A). Numbers in between brackets indicate sample sizes. Mean ± S.E.M are indicated in the graphs. Scatter-plot labelled with the same letter are not statistically different. *n.s*.: non-significant; *:p<0.05; **:p<0.01; ***:p<0.001.For full statistical details see Supplementary Table S1.

### Female modulate ejaculate ejection based on male olfactory cues

To further dissect this CFC, we explored the temporal structure of the effect of female exposure to males between first and second mating and the nature of the male cue based on which females modulate ejaculate ejection latency. We first investigated the critical time window during which CFC occurs, by subdividing the 120 minute window after the end of mating (AEM) during which sperm is stored by females (Manier et al., 2010). We exposed single mated females to a single *CS* male in the arena 30, 60, 90, or 120 minutes AEM, or kept the female isolated as a control, and recorded ejaculate ejection latency. We found that females must be exposed to *CS* males during the first 60 minutes AEM to show a decrease in ejaculate latency (Fig. 3A). However, being in the presence of a male only during the first hour AEM is not sufficient to observe a complete advancement of the timing of ejaculate ejection (Fig 3B). Together, these results suggest that modulation of ejaculate ejection latency requires continuous presence of males after the end of a female’s first mating.

**Figure 3:**
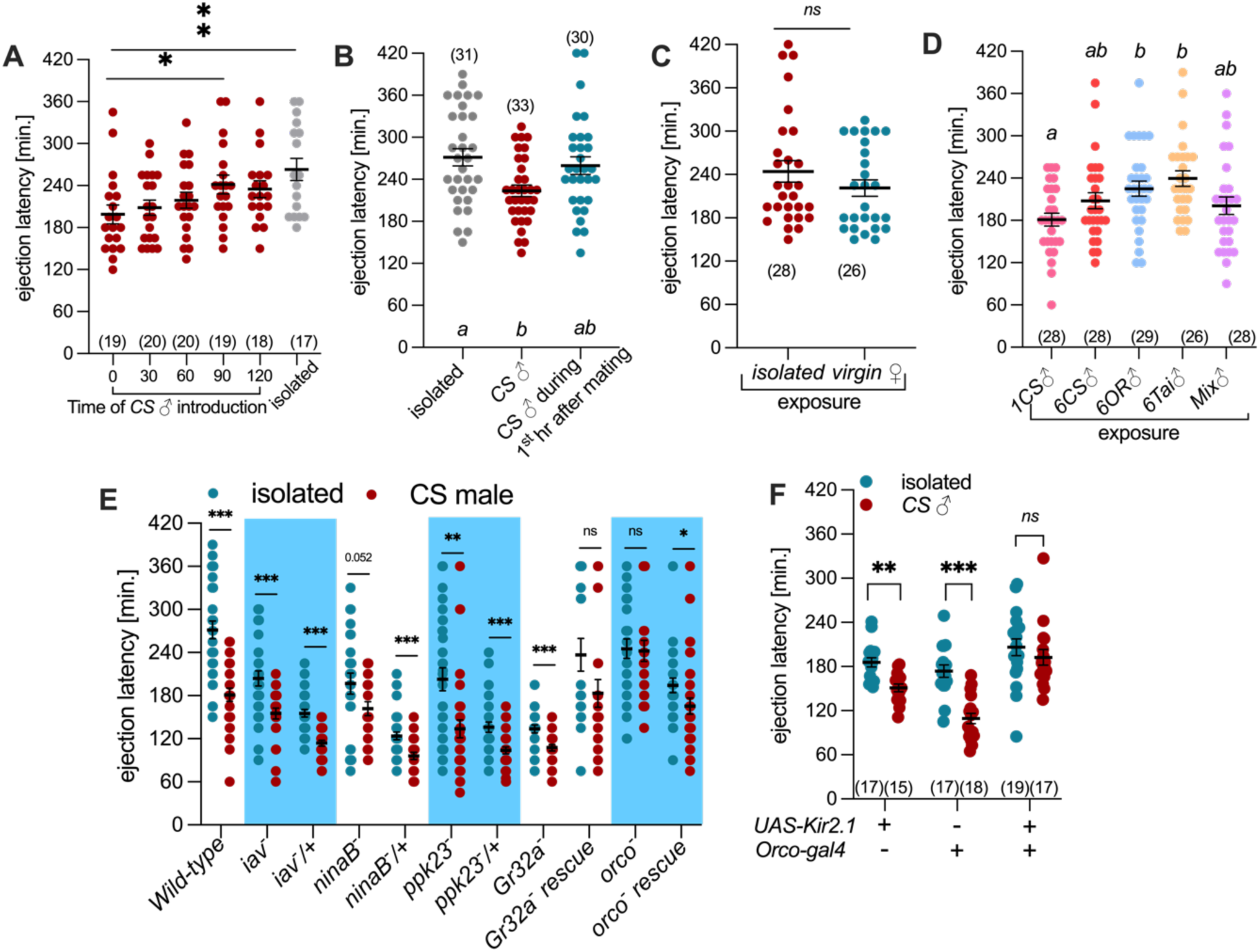
Females sense high quality males via olfaction. (A) Mean time of ejaculate ejection of a recently mated female exposed to a *CS* male, indicated by “o” or at the indicated times after she finished mating. Males were kept in the arena until ejection was observed. The “isolated” condition indicates a female that was not exposed to males. (B) Ejection latency of a females who was exposed for 1 hour with a *CS* male, after which the male was removed and the female transferred to a clean arena. (C) Mean ejection latency of a recently mated female exposed to a virgin female or left isolated. (D) Ejection latency of a recently mated females exposed to males from different strains. Mix indicates 2 *OR*, 2 *CS* and 2 *Tai* males. Experiments are performed as in Fig 1B. (E) Ejection latency of mutant females of the indicted genotypes, exposed to either a *CS* male directly after the end of their first mating or kept in isolation. (F) Females in which *Orco*-expressing neurons are silenced by means of Kir2.1 expression, and their controls, exposed to either a CS male directly after the end of their first mating or kept in isolation. Numbers in between brackets indicate sample sizes. mean ± S.E.M are indicated in the graphs. Scatter-plot labelled with the same letter are not statistically different. *n.s.:* non-significant; *:p<0.05; **:p<0.01; ***:p<0.001.For full statistical details see Supplementary Table S1.

Secondly, to investigate the cue responsible for modulation of ejaculate ejection, we hypothesized that advanced ejaculate ejection latency should be a specific to the presence of preferred males. Consistent with this, we found that ejection latency is not modulated in presence of virgin females (Fig 3C), indicating that male-specific cues are involved. To test if females modulate ejection latency based on male identity, quantity or diversity, we measured ejection latency of females placed in the presence of one or six *CS* males, or with males of different strains or a mix of two males of each strain (Fig 3D). We found that the greater number or diversity of males did not affect female ejaculate ejection latency, narrowing the relevant cue down to male attractiveness.

To determine how females sense male attractiveness cues, we measured ejection latency of females’ mutant for different sensory modalities in the presence or absence of a *CS* male (Fig 3E). Females with a mutation in *inactive*, affecting hearing*, ninaB*, affecting vision*, GR32a*, a gustatory receptor for a male pheromone, and *ppk23*, an ion channel associated with male pheromone sensing (Thistle et al., 2012), all showed a wild type response; *i.e.*, they reduced ejaculate ejection latency in the presence of a *CS* male compare to isolated controls. However, *Orco* mutant females, who have partially impaired olfaction, did not (Fig 3E). We confirmed this result by silencing *Orco* neurons using *Kir2.1* (Fig 3F) and found that females of both control groups showed early ejection in the presence of *CS* males compared to isolated controls, while the females with silenced *Orco* neurons did not. Taken together, these results suggest that females modulate timing of ejection in response to male pheromones perceived through the olfactory system.

### Females modulate timing of ejaculate ejection in response to a mix of heptanal and cVA

To test the influence of male pheromones on the modulation of ejaculate ejection latency, we generated males that do not produce cuticular hydrocarbons (CHCs), the main *Drosophila* pheromones, by genetically ablating the oenocytes (*oe^-^*), the cells that synthesize CHCs (Billeter et al., 2009). We exposed recently mated females to *oe^-^* males and observed that females did not advance ejection, as they do when exposed to a *CS* or a control male of the same genotype as *oe^-^* but without the ablation. This indicates that CHCs produced by the oenocytes contain one or more of the male cues used by females to modulate ejaculate ejection timing (Fig. 4A). Given that the most prominent male CHC, 7-Tricosene (7-T), is also expressed in higher quantity in *CS* than *OR* (Jallon, 1984), we tested the hypothesis that 7-T is the cue that females use for CFC by applying 7-T on *oe^-^* males and testing whether this resulted in a faster ejaculate ejection in females exposed to treated than in untreated *oe^-^* males. We measured ejection latency of females in the presence of *oe*^-^ males perfumed with 1000ng of 7-T, which is an average amount in *CS* males, and observed a reduction in female ejaculate ejection latency similar to that observed when females are exposed to *CS* males (Fig. 4A). Recently, we reported that 7-T is oxidized into a smaller molecule, heptanal, which is sensed by the olfactory system and is used by females to choose egg-laying sites occupied by conspecifics (Verschut et al., 2023). To determine whether 7-T, heptanal or both are sensed by females to modulate ejaculate ejection latency, we perfumed *oe*^-^ male with 1000ng of heptanal (Fig. 4A) and found that females show a decreased ejection latency in the presence of the heptanal perfumed males compared to controls, recapitulating the “high quality male”-exposed phenotype.

**Figure 4:**
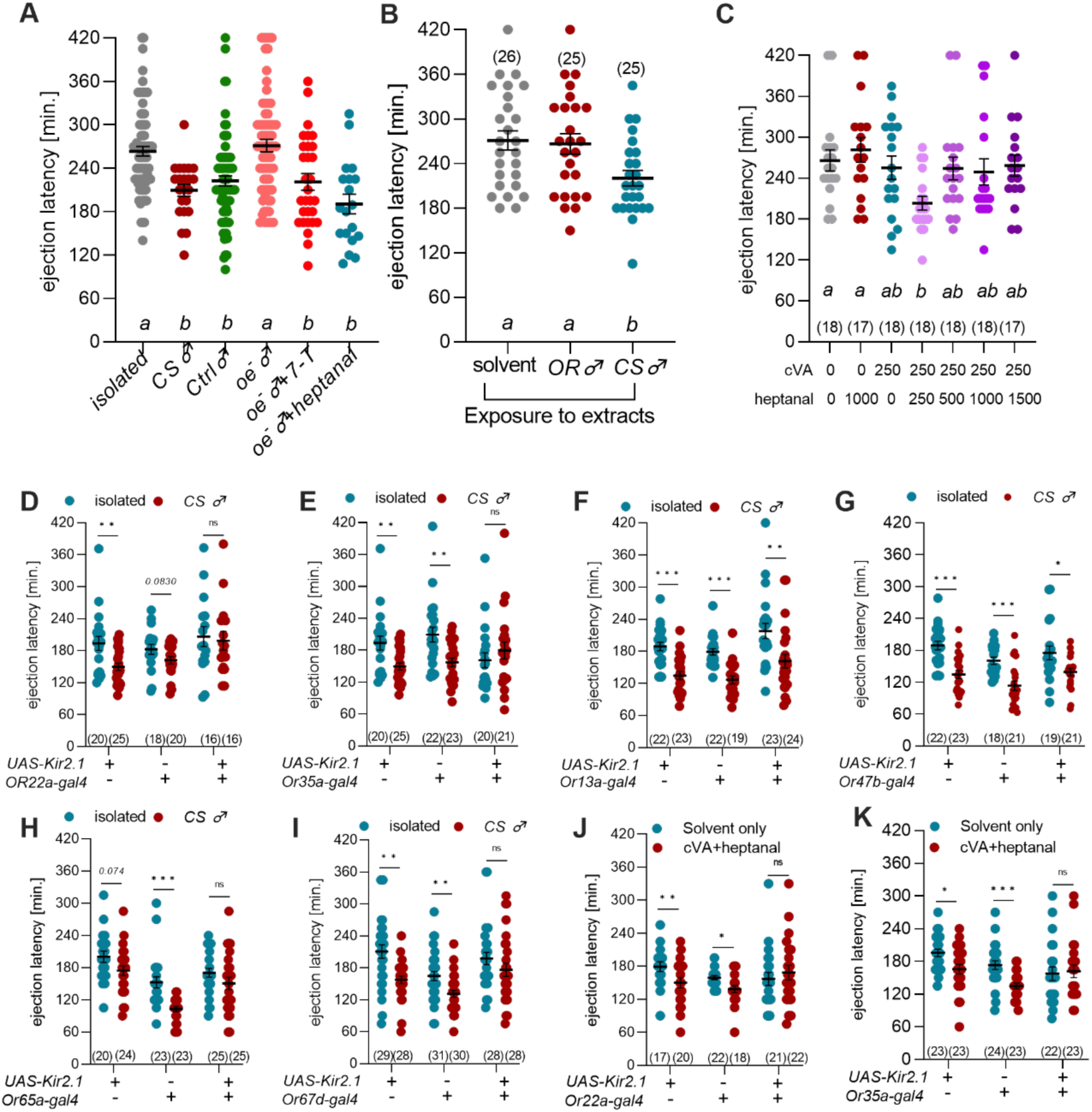
Females modulated ejection latency based on heptanal and cVA. (B) mean ejection latency of a recently mated female exposed to extracts from the equivalent of one OR or one CS male. (A) Mean ejection latency of a recently mated female exposed to males of the indicated genotypes. Genotype with “+ 7-T” where perfumed with 1000ng of this compound. Genotype with “+heptanal” were perfumed with 10000 ng of this compound. (C) mean ejection latency of a recently mated female exposed to cVA and heptanal at a variety of concentrations. (D-I) Females in which Specific Odorant receptors neurons are silenced by means of Kir2.1 expression, and their controls, exposed to either a CS male directly after the end of their first mating or kept in isolation. (J-K) same as D-I but with the males replaced by a blotting paper containing 250ng of heptanal and 250ng of cVA. Numbers in between brackets indicate sample sizes. mean ± S.E.M are indicated in the graphs. Scatter-plot labelled with the same letter are not statistically different. *n.s*.: non-significant; *:p<0.05; **:p<0.01; ***:p<0.001.For full statistical details see Supplementary Table S1.

Because flies deposit pheromonal residues in their environment (Duménil et al., 2016; Keesey et al., 2017; Verschut et al., 2023), we next tested whether male pheromones alone can modulate female ejaculate ejection without the male directly interacting with the females. For this, we exposed mated females to pheromonal extracts of *CS*, *OR* males or a solvent control, placed on a filter paper in a cup covered with a mesh, and measured the timing of ejection of females exposed to these extracts between their first and second mating. We observed reduced ejection latencies of females in the presence of *CS* pheromone extracts compared to *OR* pheromone extracts or a hexane control, demonstrating that male pheromones are sufficient to modulate female ejaculate ejection latency (Fig. 4B). To isolate the specific pheromone responsible for this effect, females were exposed to individual pheromones in a dose response manner. Females exposed to cVA, 7-T or heptanal alone did not advance timing of ejection at any doses (Supp Fig. 4A,B,C). As cVA acts in combination with other pheromones and is still expressed in oe^-^ males (Billeter et al., 2009; Laturney & Billeter, 2016; Verschut et al., 2023), we surmised that females respond to heptanal perfumed *oe^-^*because of the resulting combination of cVA and heptanal. To test this hypothesis, we exposed females to a mix of heptanal and cVA. Females indeed advanced ejaculate at specific doses of heptanal and cVA demonstrating that the cue indicative of male quality is a combination of cVA and heptanal (Fig 4C).

Next, we investigated which odorant receptor neurons (ORN) females use to sense male quality/cVA-heptanal and adjust ejaculate ejection in response. We silenced different ORNs by driving expression of *UAS-Kir2.1* with ORN specific Gal4 driver lines. *ORN22a*, *ORN13a*, and *ORN35a* were tested because they are necessary for females to respond to heptanal in the context of egg-laying site choice (Verschut et al., 2023). Silencing ORN22a+ (Fig4.D) and ORN35a+ (Fig 4E) neurons resulted in females no longer advancing timing of ejaculate ejection in the presence of *CS* males. However, we observed advanced ejection latency in the presence of *CS* males when ORN13a neurons were silenced (Fig 4F). Together, these data demonstrate that ORN22a and ORN 35a are the main inputs for the modulation of ejaculate ejection in response to heptanal. As an out-group control, we also tested ORN47b, which we showed to be involved in mated female mate choice in response to palmitoleic acid (Kohlmeier et al., 2021), but this ORN does not appear necessary for the modulation of ejaculate ejection (Fig 4G). We next tested the two documented receptors for cVA, *ORN65a* and *ORN67d* (van der Goes van Naters & Carlson, 2007). Females with silenced ORN65a (Fig4.H) or ORN67d (Fig 4I) neurons did not significantly decrease ejection latency in the presence of a *CS* male, indicating that they are responsible for sensing cVA in the context of ejaculate ejection modulation. We confirm the involvement of OR22a and OR35a (Fig 4J,K) by exposing mutant and control females to a mix of heptanal and cVA inducing a decrease of the timing of ejaculate ejection in control flies. In both cases females with silenced ORN22a 35a neurons did not advance timing of ejection. We conclude that female adjust ejaculate ejection latency in response to preferred males through a combination of heptanal receptors OR22a and OR35a and cVA receptors OR65a and OR67d.

## Discussion

We provide the first demonstration, to our knowledge, of a mechanism for cryptic female choice in response to male cues in *D. melanogaster*. We show that *D. melanogaster* females advance timing of ejaculate ejection when exposed to a preferred male after their first mating, leading to decreased sperm stored from the first male and an increase in paternity share of their second mate. Females assess male quality based on olfaction, sensing two male pheromones, cVA and heptanal, an oxidization product of 7-T. Our results highlight that social cues happening outside of mating itself can influence paternity.

### Females exert male selection at multiple stages of reproduction in *D. melanogaster*

We recently showed that virgin *D. melanogaster* females mate less selectively as virgins than as mated females (Kohlmeier et al., 2021). Once females have secured sperm from their first mating, they become more selective and will mate with higher quality males if they encounter them (Kokko & Mappes, 2005). Because of last male sperm precedence, the second male will sire most of the offspring, reducing paternity from the first male. The cryptic female choice mechanism we report here further extends the importance of post-copulatory events by showing that females can influence paternity by modulating ejaculate ejection timing and biasing the ratio of sperms they keep from their first mate depending on their sesning of higher quality males in their environment. Critically, we show that males do not need to be physically present as exposure to specific male pheromones in the physical absence of a male is sufficient to modulate ejaculate ejection timing. Given that *D. melanogaster* naturally deposit pheromones where they dwell (Duménil et al., 2016; Keesey et al., 2017; Verschut et al., 2023), we speculate that environmental pheromones allow recently mated females to determine the presence of high-quality males without having to physically interact with them. Indeed, those recently mated females are in the process of storing sperm from their first mate and are not yet attractive to males because they have also acquired anti-aphrodisiac pheromones from their first mate (Laturney & Billeter, 2016; Manier et al., 2010). These anti-aphrodisiac pheromones are jettisoned by the female together with excess ejaculate once the sperm storage process is over; a point at which female may begin remating (Laturney & Billeter, 2016). Advancement of ejaculate ejection in presence of preferred males may thus allow females to allocate a higher ratio of paternity when they eventually remate with those males.

A picture is emerging where *D. melanogaster* female adapt reproductive behaviors to match their social environment. Indeed, by mating with low discrimination as a virgin, they ensure they acquire sperm as soon as possible. By adjusting how much sperm they keep from the first male, they can adjust investment in that first male depending on their perception of mating opportunities with higher quality males. *D. melanogaster* female have thus multiple levels of controls over who sires their offspring, allowing them to maximize fitness in different demographic conditions; relaxing sexual selection when male quality is low and making it more stringent when male quality is high.

### Female advance ejaculate ejection when sensing specific male pheromones

Our findings reveal that female advance ejaculate ejection in response to the sensing of male pheromones cVA and heptanal, a derivative of 7-Tricosene (Verschut et al., 2023). cVA and 7-T are male pheromones that function together to repel other males; they are naturally expressed by males and are part of an anti-aphrodisiac cocktail transferred by males to recently mated females, which renders them unattractive (Billeter et al., 2009; Laturney & Billeter, 2016). In this context, males sense 7-T via Gustatory receptor Gr32 expressed in their legs that get activated when touching mated females, while cVA is detected via their olfactory system (Billeter & Levine, 2013; Laturney & Billeter, 2016; Wang et al., 2011). We show here that females also use this pheromonal combination as a cue to modulate ejaculate ejection latency. However, they do not sense 7-T via taste receptors, but its breakdown product heptanal via olfaction. This indirect detection of 7-T via olfaction in females as opposed to direct detection by gustatory receptor in males, allows 7-T to take a different function in males and females. Detection by olfaction is typically at a distance, allowing females to sense males without physically interacting with them. That females use olfactory cues rather than gustatory ones requiring physical contact, gives support to our proposal that females advance ejaculate ejection in response to their evaluation of males they will potentially mate with in the future, as opposed to those they are in direct contact with.

A second surprising aspect of the discovery that cVA and heptanal modulate ejaculate ejection is that these pheromones have not been implicated as major contributors to female mate choice, at least not for mated females. Indeed, the change in female mate selectivity between virgin and mated females is mediated by a different pathway involving odorant receptor neurons Or47b and male pheromone palmitoleic acid (Kohlmeier et al., 2021). Our results indicate that this pathway is not involved in CFC as silencing Or47b ORN in females still results in advanced ejaculate ejection in the presence of high-quality males. Why would females have evolved different pathways for mate selection and ejaculate ejection? Mate choice requires close temporal and spatial association between the cue and a mate, while our data show that ejaculate ejection is a slow build up requiring long exposure to male cues. These spatio-temporal differences in decision making might have resulted in the evolution of two separate pathways, one for evaluating present mating opportunities and one for evaluating future mating opportunities. Alternatively, the cues themselves, palmitoleic acid for mate choice and cVA/heptanal for ejaculation ejection, may have different volatility and/or stability, as well as spatial distributions. A potential lower volatility/stability, or a lack of deposition of palmitoleic acid in the environment would prevent it from exerting an environmental effect and would thus remain a cue physically associated with a male. cVA and 7-T are however documented to be deposited in the environment, and are both detectable at a distance.

### Females bias paternity through sperm allocation in the seminal receptacle

While we do not provide the full pathway linking sensing of male cues as an input and changing sperm storage and cryptic female choice as an output, our data and published findings allow speculations on the nature of this pathway. The female olfactory system is the input that starts the chain of events leading to differential sperm storage in response to male exposure. Lee et al., (2015) demonstrated that female neurons expressing the Dh44 peptide function to temporally block ejaculate ejection, allowing females sufficient time to store sperm before eventually ejecting the surplus ejaculate a few hours after mating. It is likely that there exists a connection between olfactory processing and Dh44-expressing neurons. Indeed, a recent report shows that exposure to male pheromones activate pC1 neurons, neurons associated with female receptivity (Zhou et al., 2014), in the female brains, resulting in faster ejaculate ejection and remating (Yun et al., 2023). If there is a connection between pC1 neurons and Dh44 neurons, detection of pheromones indicative of the presence of a preferred male by the olfactory system may activate pC1 neurons, who would then reduce inhibition of ejaculate ejection by Dh44 neurons resulting in advancement of ejaculate ejection. In absence of high-quality males, pC1 neurons may be less activated and fail to increase Dh44 neurons activity resulting in a default ejection a few hours after mating, ensuring maximum storage of sperm from their first mate. How would these central neurons then influence ejaculate ejection? While there is no direct connection between Dh44 neurons and the female reproductive tract, the two sperm storage sites are innervated by descending neurons from the brain, including octopaminergic/thyraminergic neurons (Avila et al., 2012). Interestingly, blocking functioning of these neurons impairs sperm storage (Avila et al., 2012), and can even bias sperm use in twice mated females (Chen et al., 2022). This strongly suggest that these neurons are cryptic female choice effectors, who might receive input from neurons integrating male pheromones sensing by pC1 neurons and timing by Dh44 neurons. Future work will doubtless elucidate the neuronal circuit integrating male cues with sperm storage and ejaculate ejection latency.

## Material and Methods

### *Drosophila* strains and Rearing conditions

*Drosophila melanogaster* strains were reared in incubators set at 25°C with a 12:12h light–dark cycle (LD 12:12) on food medium containing agar (10 g l−1), glucose (167 mM), sucrose (44 mM), yeast (35 g l−1), cornmeal (15 g l−1), wheatgerm(10 g l−1), soya flour (10g l−1), molasses (30 g l−1), propionic acid (5 ml of 1 M) and Tegosept (2 g in 10 ml of ethanol). Virgin adults were collected using CO_2_ anesthesia and aged in same-sex groups of 15 in food vials (25×95 mm) containing food medium (23×15 mm) for 5 to 7 days before testing.

GFP and RFP males were obtained by backcrossing the original line with *Canton-S* or *Oregon-R* strains for 18 generations.

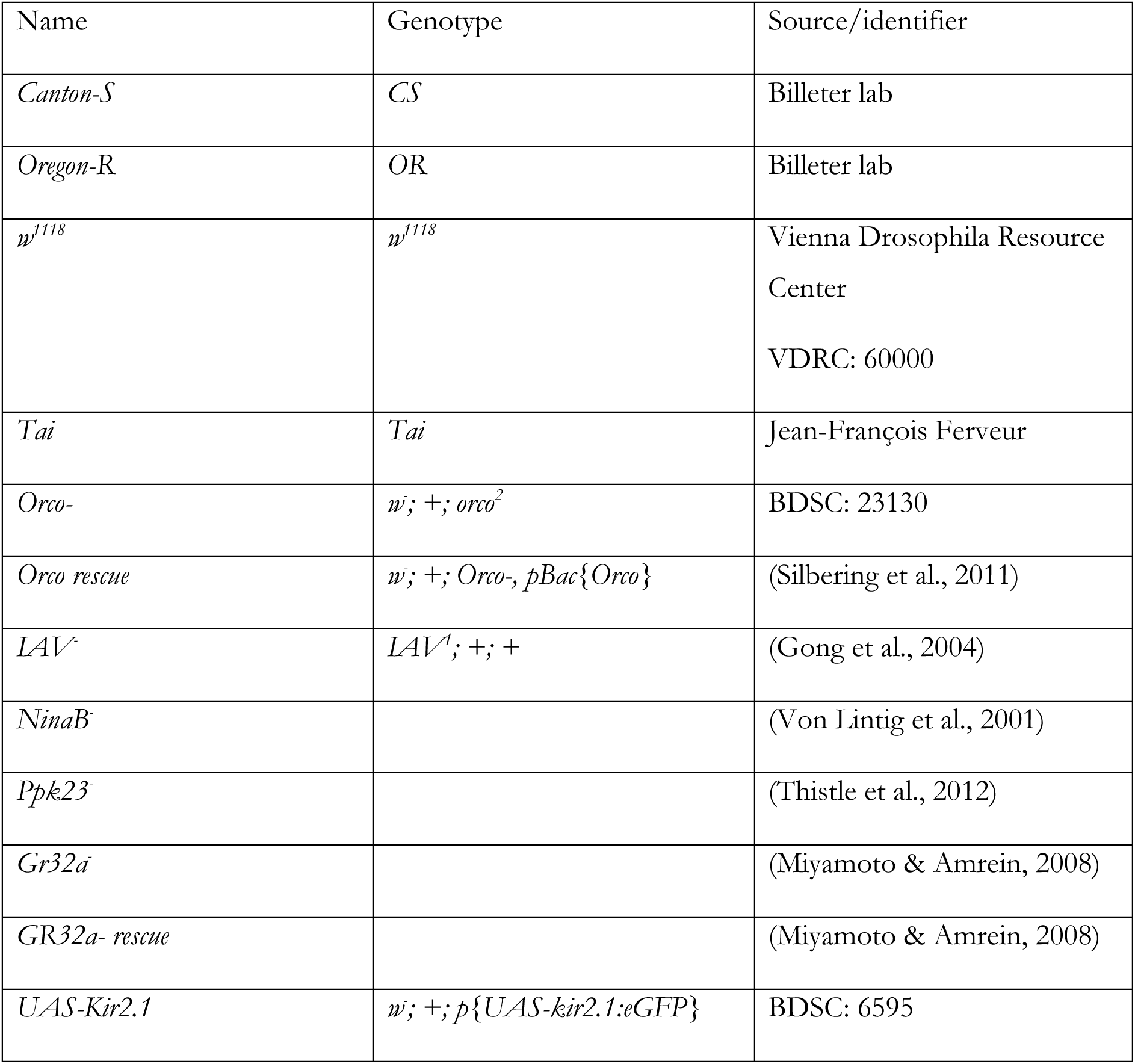
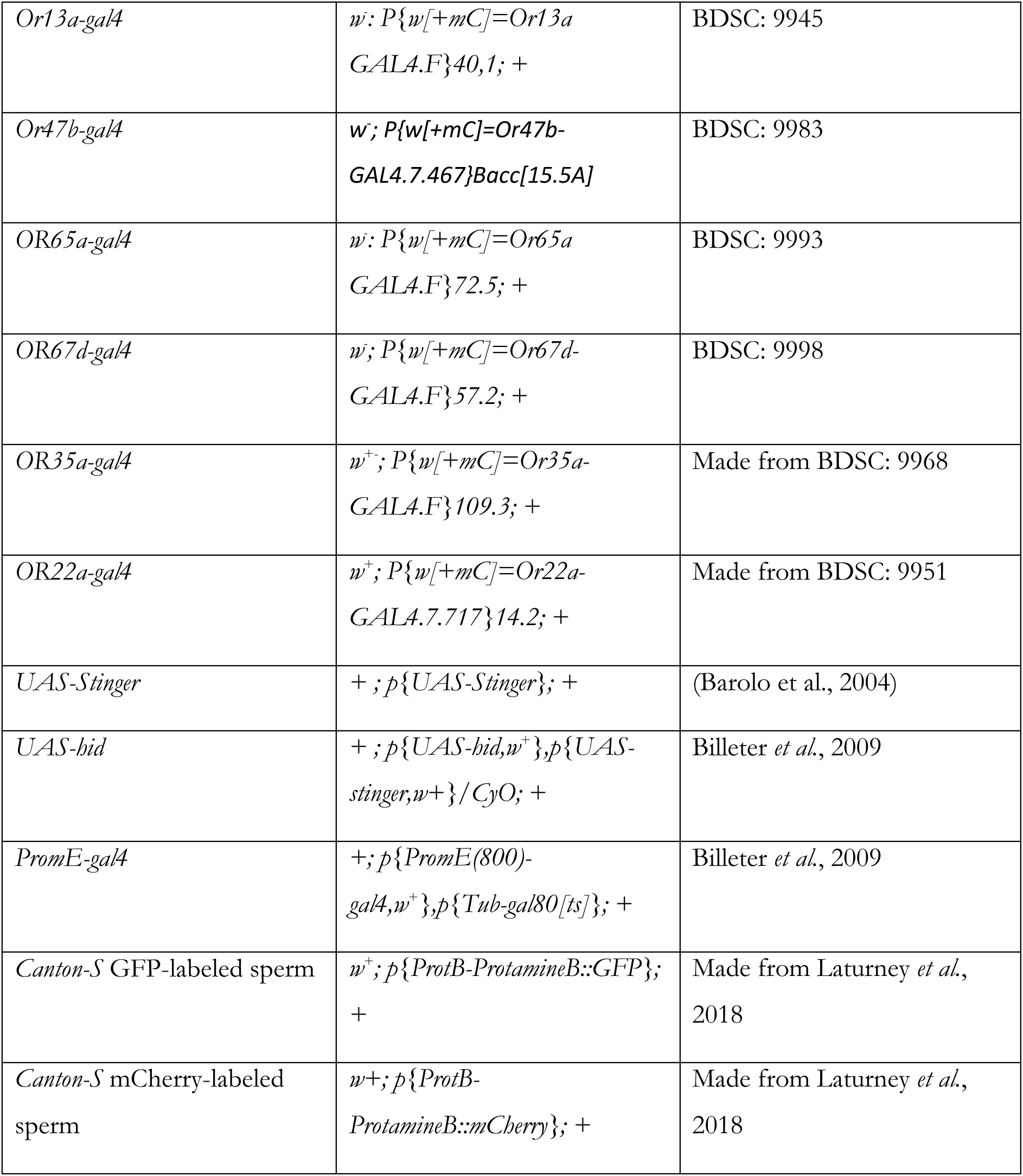

### Mate-choice assay

Mate-choice assays were performed as detailed in (Kohlmeier et al., 2021; Verschut et al., 2023). Briefly, coloring vials are made by putting 0.5g of green or pink fluorescent powder (Slice of the moon) in an empty vial and rotating it until the surface gets covered. The residual powder was removed by tapping down the vial. 15 males of both strains were transferred in a different coloring vial, the vial was rotated for 3s. A male of each of the two strains are then individually transferred in a mating arena for 30min to recover. Colors were randomly assigned on every experimental day. Females are then individually added to the arena and inspected for mating every 10min for 60min. The flies that did not mate within 60min were discarded.

### Ejaculate ejection timing assay

*Canton-S* virgin female (or different genotype based on experiment) were individually mated at 25°C with a single *Canton-S* (CS) male in a mating arena (35 x 10mm Petri dish, Greiner Bio-One, containing 2mL of fly food) at ZT1 under light condition. After mating, females were checked with a UV flashlight (Perel EFL41UV) to ensure the presence of the mating plug indicative of matin (Lung & Wolfner, 2001). Females that did not mate within 2 hours were discarded; that corresponded to less than 10% of trials. Mated females were transferred individually using a mouth pipette in a fresh mating arena alone, with a new male or pheromones. Ejection of the mating plug was checked every 15min with a UV flashlight for 7h after the end of mating by checking its presence in the arena and on the female genitals. Experiments were performed in a temperature-controlled chamber at 25°C on a bench cleaned daily with ethanol/water and 100% Acetone to avoid accumulation of pheromonal residues.

### Twice-mated female assay

The beginning is identical to the ejaculate ejection timing assay. After ejection, females were transferred in a new mating arena with a GFP-labeled sperm male in *Canton-S* background. Remating with the new male was visually inspected every 10min for 4 hours. Females that did not remate after 4 hours were discarded; that corresponded to less than 20% of trials. After remating, the presence of the new mating plug was checked and the female was transferred alone to a new mating arena until ejection of the 2^nd^ mating plug. Ejection of the mating plug was checked every 15min. After the 2^nd^ ejection, females were individually transferred in a fresh food vial every 2 days for 7 days. Offspring coming from these vials were counted and their paternity assessed as in Laturney *et al*. (2018) by the red fluorescence in the ocelli due to *M{3xP3-RFP.attP}ZH-102D* insertion when they are fathered by the male from GFP-labeled sperm lines using a fluorescent stereo microscope (Leica MZ10 F). For sperm counting experiments, females followed the same protocol but were first mated with a mCherry-labeled sperm male backcrossed in a *Canton-S* background. After the 2^nd^ ejection they were placed individually in a 0.5mL Eppendorf tube and were immediately flash-frozen in liquid nitrogen directly after ejection and stored at −20°C until dissection. The fluorescent-labeled sperm male in *Canton-S* background were obtain by backcrossing the original lines to *Canton -S* for 18 generations.

### Sperm count

Flash frozen female reproductive tract was dissected in PBS 1X and mounted on Vectashield (Vector Laboratories) on a glass slide covered with a coverslip elevated by bridging it in between two coverslips. The reproductive tracts were imaged with a Zeiss LSM 800 confocal microscope with a 40x water immersion lens using a 587nm excitation wavelength for mCherry and 488nm for GFP. Sperm are manually counted using Fiji software as in Laturney et al., (2018). Counts were done blind, each female being given a random ID, and was a recounted blind again by another experimenter.

#### Pheromonal exposure

*CS* and *OR* males were anesthetized on ice in a plastic vial then transferred using forceps cleaned by dipping them in n-hexane, methanol and n-hexane again in a 2ml glass vial. 10µL of hexane/flies are pipetted in the glass vial for the extraction. The vial is vortexed for 3 minutes, every minute the vial is checked to ensure that no flies are stuck above the hexane. The supernatant is pipetted in a clean 2mL glass vial.

Filter paper discs were made using a 3 mm diameter one-hole paper puncher out of cellulose chromatography paper rolls. Using clean forceps, the discs were placed in a well, made of the cap of 0.5mL reaction tube (Sarstedt). 10µL of pheromone solution were pipetted on the filter paper and the well was put under a nitrogen flow for 5min to let the hexane evaporate. The wells were then covered with foil paper, to avoid direct contact of flies with the pheromone patch, and placed at the center of mating arena. 2mL food was poured around the edges of the well make them stick to the arena. Seven holes were made in the foil paper, small enough to not let a fly go through and let the odour spread in the arena just before introducing a mated female inside.

#### Perfuming

Oenocyte ablated flies were generated by crossing *+; PromE(800)-Gal4 [2M],Tub:Gal80^ts^; +* to +: *UAS-StingerII, UAS-hid/CyO; +* flies (Billeter et al., 2009). Flies are reared at 18°C and transfer at 25°C after the adults have been collected for at least 5 days to allow the expression of the Gal4 and the ablation of the oenocytes. The pheromone of interest (or hexane for control) at 28 times the desired concentration (7 flies are perfumed at the same time with a 25% transfer efficiency) per fly is pipetted in a 2ml glass vial and briefly vortexed. The vial is placed under a nitrogen flow until complete evaporation of the hexane. As in the CHCs extraction, flies are anesthetized on ice, 7 of them are transferred by their wings into the coated glass vial with clean forceps. The vial is vortexed at low speed 3 times for 20s with a 20s interval. The flies are transferred into a fresh food vial and an hour of recovery time is given to them before the start of the experiment.

#### Gas chromatography

For hydrocarbon analysis, single flies were placed in glass microvials containing 50 μl of an internal standard solution made by dissolving 10 ng/ml of octadecane (C18) and 10 ng/ml of hexacosane (C26) in hexane. The vials were vortexed at minimum speed for 2min and flies were removed using a clean metal pin. The resulting hydrocarbon extracts were analysed using an Agilent 7890 gas chromatograph with a flame ionization detector, an Agilent DB-1 column (Diameter: 0.180 mm; film 0.18 μm) and a split-splitless injector set at 250°C with 40ml/min splitless flow. Injector valve opened 1.5 min after injection in splitless mode with helium as carrier gas (flow: 37.2 cm/sec). The oven temperature program begins at 50 °C for 1.5 min, ramping at 10°C /min to 150 °C, then ramping at 4 °C /min to 280°C and holding for 5 minutes. ChemStation software (Agilent technologies) was used to quantify compounds based on peak areas relative to internal standard C26 as described in Verschut et al., 2022.

### Statistics

Statistical analyses were performed using R 4.3.0 software (The R Foundation for Statistical Computing). Residuals were visually inspected for normality, and homogeneity of variances were evaluated with the F-test. Some data were log transformed when normality was not respected (see supp table). Wilcoxon-Mann-Whitney test and Student’s t-test were used to compare ejection latency, proportion of offspring, number of offspring and number of sperm between two groups (isolated vs *CS* male or *CS* male vs *OR* male). Kruskal-Wallis test followed by Dunn’s multiple comparisons test and the Anova test followed by Tukey post-hoc comparisons or Dunnett’s test were used to compare ejection latency between more than two groups.

All graphs and some statistical analyses were performed using GraphPad Prism 9.5.1 software.

## Acknowledgment

We thank Meghan Laturney for helpful feedback and comments on the manuscript. We thank Thomas Verschut for his help with the pheromones and how to handle them. Thanks to Gerard Overkamp for helping to solve technical issues with the experimental work. We thank Simon Robledo-Cardona for helping setting up pilots for some these experiments. This work was funded by the Netherlands Organization for Scientific Research (NWO) ALWOP.611 to J.C. Billeter and by a fellowship of the German Research Foundation awarded to P.K. (KO 5835/2-1).

---

**Supplementary table 1.**

| Fig | Test | Response variable | Explanatory factor | Results | Post-hoc test (only p-value <0.05) |
| --- | --- | --- | --- | --- | --- |
| 1B | Anova | Ejection latency | Post-mating condition [male CS, male OR, isolated] | F=10.801, df=2, p=5.82e-05 | Male CS-isolated: p=0.0002243, Male OR-isolated: p=0.9683661, Male OR- Male CS: p= 0.0004802 |
| 1C | Unpaired t-test | Proportion of offspring | Post-mating condition [male CS, male OR] | t = -2.0914, df = 93, p-value = 0.03922 | NA |
| 1D | Unpaired t-test | Number of offspring | Post-mating condition [male CS, male OR] | t = -2.0084, df = 93, p-value = 0.0475 | NA |
| 1E | Mann-Whitney exact test | Number of offspring from the 1<sup>st</sup> male | Post-mating condition [male CS, male OR] | U = 817.5, p-value = 0.0204, two-tailed | NA |
| 1F | Unpaired t-test | Number of offspring from the 2nd male | Post-mating condition [male CS, male OR] | t = 0.0092772, df = 93, p-value = 0.9926 | NA |
| 2B | Mann-Whitney exact test | Proportion of offspring in SR | Post-mating condition [male CS, male OR] | U = 62, p= 0,0039, two-tailed | NA |
| 2C | Mann-Whitney exact test | Number of sperm in SR | Post-mating condition [male CS, male OR] | U = 114,5, p= 0,3170, two-tailed | NA |
| 2D | Unpaired t-test | Number of sperm in SR first male | Post-mating condition [male CS, male OR] | t = -2.3157, df = 32, p-value = 0.02714 | NA |
| 2E | Unpaired t-test | Number of sperm in SR 2nd male | Post-mating condition [male CS, male OR] | t = 2.9467, df = 32, p-value = 0.005949 | NA |
| 2F | Mann-Whitney exact test | Proportion of offspring in SP | Post-mating condition [male CS, male OR] | U= 85, p=0,0715, Two-tailed | NA |
| 2G | Unpaired t-test | Number of sperm in SP | Post-mating condition [male CS, male OR] | t=0,4014, df=32, p-value=0,6908 | NA |
| 2H | Unpaired t-test | Number of sperm in SP first male | Post-mating condition [male CS, male OR] | t = -1.6899, df = 32, p-value = 0.1008 | NA |
| 2G | Unpaired t-test | Number of sperm in SP 2nd male | Post-mating condition [male CS, male OR] | t = 0.54826, df = 32, p-value = 0.5873 | NA |
| 3A | ANOVA | LOG transformed ejection latency | Time of male entrance [0min, 30min, 60min, 90min, 120min, isolated] | Df=5, F= 3.1999, p-value = 0.00994 | Dunnett's multiple comparisons test:<br>0 vs isolated: p=0,0025<br>0 vs. 30: p=0.7141<br>0 vs. 60: p=0.5032<br>0 vs. 90: p=0.0423<br>0 vs. 120: p=0.0914 |
| 3B | ANOVA | Ejection latency | Post-mating condition [male CS, isolated, maleCS for 1<sup>st</sup> hour AEM] | F= 5.1655, df=2, p= 0.007501 | TukeyHSD:<br>Alone-1hrCSmale: p=0.7335201<br>Csmale-1hrCSmale: p=0.0623864<br>Csmale-Alone: p=0.0076787 |
| 3C | Mann-Whitney exact test | Ejection latency | Post-mating condition [virgin female CS, isolated] | U =303, p-value = 0.2932, two-tailed | NA |
| 3D | ANOVA | Log transformed ejection latency | Post-mating condition [1male CS, 6malesCS, 6males OR, 6males tai, mix] | F=4,451, df=4, p=0,0021 | TukeyHSD:<br>1CS♂ vs. 6OR♂: p=0,0261<br>1CS♂ vs. 6Tai♂: p=0,0018 |
| 3E wild type | Unpaired t test | Ejection latency | Post-mating condition [male CS, isolated] | t=5,789, df=57, p-value<0,0001 | NA |
| 3E iav- | Mann-Whitney exact test | Ejection latency | Post-mating condition [male CS, isolated] | U=179,5, p-value=0,0012, two-tailed | NA |
| 3E iav/+ | Mann-Whitney exact test | Ejection latency | Post-mating condition [male CS, isolated] | U=180, p-value <0,0001, two-tailed | NA |
| 3E ninaB- | Unpaired t test with Welch's correction | Ejection latency | Post-mating condition [male CS, isolated] | t=1,993, df=41,19 p-value=0,0529 | NA |
| 3E ninaB-/+ | Mann-Whitney exact test | Ejection latency | Post-mating condition [male CS, isolated] | U=269, p-value =0,0005, two-tailed | NA |
| 3E ppk23- | Mann-Whitney exact test | Ejection latency | Post-mating condition [male CS, isolated] | U=289,5, p-value =0,0014, two-tailed | NA |
| 3E ppk23-/+ | Mann-Whitney exact test | Ejection latency | Post-mating condition [male CS, isolated] | U=202, p-value =0,0003, two-tailed | NA |
| 3E Gr32a- | Mann-Whitney exact test | Ejection latency | Post-mating condition [male CS, isolated] | U=111, p-value =0,0004, two-tailed | NA |
| 3E Gr32a-rescue | Mann-Whitney exact test | Ejection latency | Post-mating condition [male CS, isolated] | U=119.5, p-value =0,0733, two-tailed | NA |
| 3E orco- | Mann-Whitney exact test | Ejection latency | Post-mating condition [male CS, isolated] | U=305, p-value =0,7109, two-tailed | NA |
| 3E orco-rescue | Mann-Whitney exact test | Ejection latency | Post-mating condition [male CS, isolated] | U=336, p-value =0,0108, two-tailed | NA |
| 3F | Unpaired t test | Ejection latency | Post-mating condition [male CS, isolated] | Kir/+: t=4,166, df=30, p=0,0002<br>Orco/+: t=5,970, df=33, p<0,0001<br>orco/kir :t=0,8782, df=34, p=0,3860 | NA |
| 4A | ANOVA | Ejection latency | Post-mating condition [male CS CHCs, male OR CHCs, hexane] | F= 3.4526, df=2, p= 0.03749 | TukeyHSD:<br>hexane-CS CHC: p=0.0289712<br>OR CHC-CS CHC: p=0.4679140<br>OR CHC-hexane: p=0.3060676 |
| 4B | Kruskal-Wallis test | Ejection latency | Post-mating condition [isolated, male CS, male stinger, male oe-, male oe- +7T, male oe- + heptanal] | P <0.0001 | Dunn's multiple comparisons test:<br>isolated vs. CS♂: p=0,0018<br>isolated vs. Ctrl♂: p=0,0002<br>isolated vs. oe♂: p>0,9999<br>isolated vs. oe♂+7-T: p=0,0093<br>isolated vs. oe♂+heptanal: p<0,0001 |
| 4C | ANOVA | LOG transformed ejection latency | Post-mating condition [hexane, 250cVA, 1000heptanal, 250cVA+250heptanal, 250cVA+500heptanal, 250cVA+1000heptanal, 250cVA+1500heptanal] | F-value= 2.575, df=6, p-value: 0.02226 | TukeyHSD:<br>Hexane vs 250+250: p= 0.0452316<br>0+1000 vs 250+250: 0.0090598 |
| 4D | Unpaired t test | Ejection latency | Post-mating condition [male CS, isolated] | Kir/+: p=0,0044<br>OR22a/+: p=0,0830 | NA |
|  |  |  |  | OR22a/kir: p=0,7615 |  |
| 4E | Mann-Whitney exact test | Ejection latency | Post-mating condition [male CS, isolated] | Kir/+: p=0,0044<br>OR35a/+: p=0,0024<br>OR35a/kir: p=0,2679 | NA |
| 4F | Mann-Whitney exact test | Ejection latency | Post-mating condition [male CS, isolated] | Kir/+: p<0,0001<br>OR13a/+: p<0,0001<br>OR13a/kir: p=0,0016 | NA |
| 4G | Mann-Whitney exact test | Ejection latency | Post-mating condition [male CS, isolated] | Kir/+: p<0,0001<br>OR47b/+: p=0,0001<br>OR47b/kir: p=0,0176 | NA |
| 4H | Mann-Whitney exact test | Ejection latency | Post-mating condition [male CS, isolated] | Kir/+: p=0,0741<br>OR65a/+: p<0,0001<br>OR65a/kir: p=0,1592 | NA |
| 4I | Mann-Whitney exact test | Ejection latency | Post-mating condition [male CS, isolated] | Kir/+: p=0,0007<br>OR67d/+: p=0,0029<br>OR67d/kir: p=0,1518 | NA |
| 4J | Mann-Whitney exact test | Ejection latency | Post-mating condition [cVA+heptanal, hexane] | Kir/+: p=0,0044<br>OR22a/+: p=0,0063<br>OR22a/kir: p=0,5085 | NA |
| 4K | Mann-Whitney exact test | Ejection latency | Post-mating condition [cVA+heptanal, hexane] | Kir/+: p=0,0171<br>OR35a/+: p=0,0001<br>OR35a/kir: p=0,6217 | NA |
| Supp Fig1 | Mann-Whitney exact test | Ejection latency | Post-mating condition [male CS, male OR] | P=0,2553 | NA |
| Supp Fig4A | Kruskal-Wallis test | Ejection latency | Post-mating condition [hexane, cVA 100, cVA 250, cVA 500, cVA 1000] | P=0,3431 | NA |
| Supp Fig4B | Kruskal-Wallis test | Ejection latency | Post-mating condition [hexane, 7-t 100, 7-t 250, 7-t 500, 7-t 1000, 7-t 1250, 7-t 1500] | P=0,9331 | NA |
| Supp Fig4C | Kruskal-Wallis test | Ejection latency | Post-mating condition [hexane, heptanal 200, heptanal 500, heptanal 1000, heptanal 1250, heptanal 1500] | P=0,2531 | NA |

## Supplementary figures

**Supplementary figure 1:**
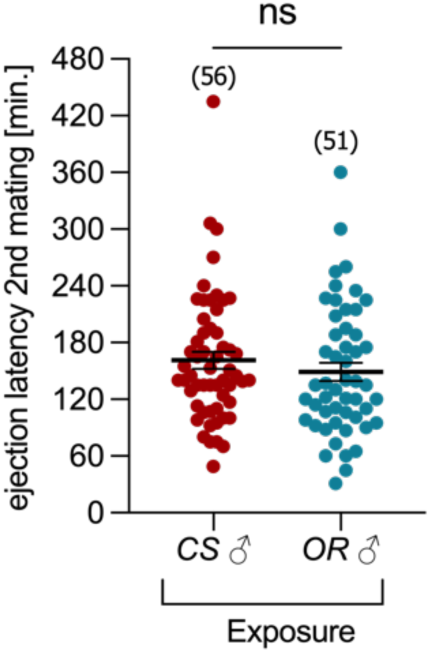
Female 2^nd^ male ejaculate ejection latency is not affected by exposure to different types of males. Ejaculate ejection latency of females after a second mating exposed to the presence of *CS* or *OR* males between matings. Numbers in between brackets indicate sample sizes. mean ± S.E.M are indicated in the graphs. Scatter-plot labelled with the same letter are not statistically different. *n.s*.: non-significant; For full statistical details see Supplementary Table S1.

**Supplementary figure 4:**
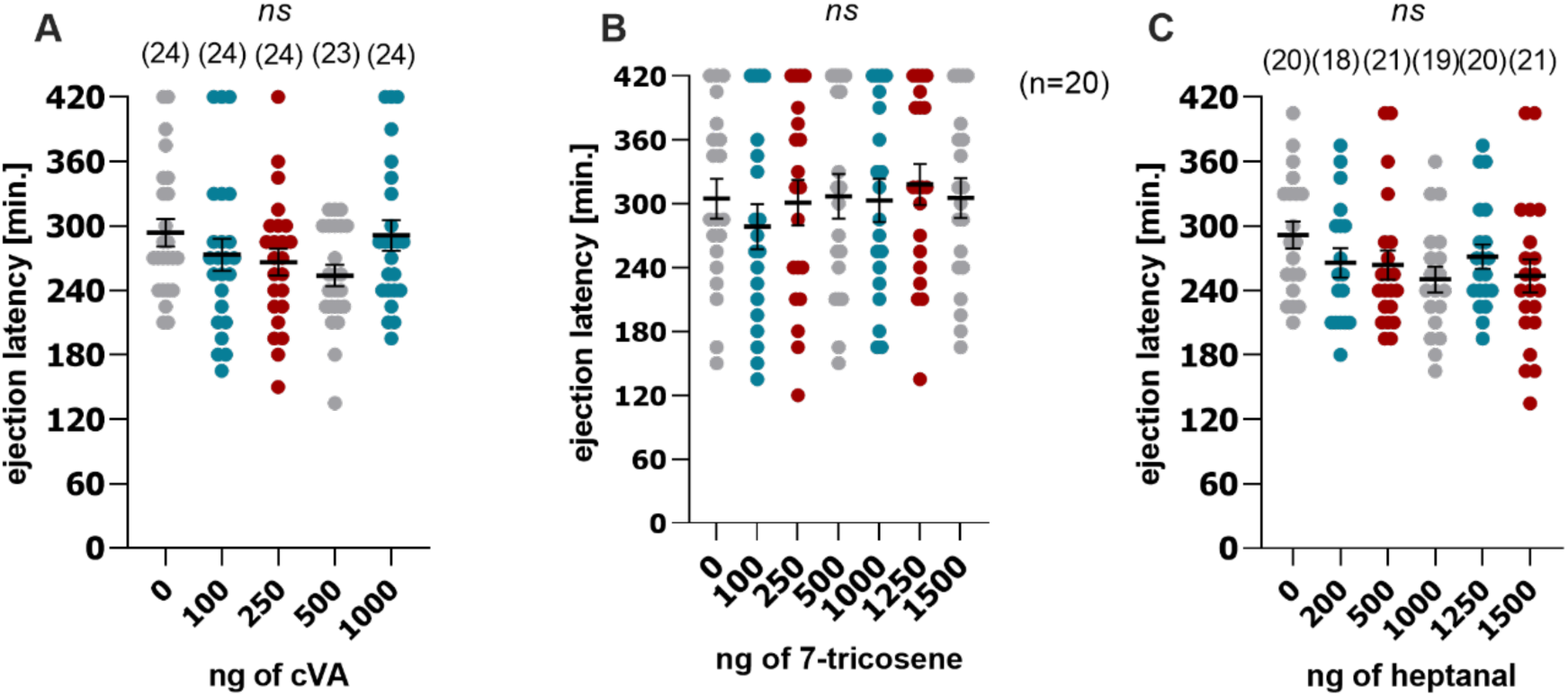
Ejection latency in response to male’s pheromones. (A) Ejaculate ejection latency of female *CS* in response to different amount of cVA or to the solvent only (hexane). (B) Ejaculate ejection latency of female *CS* in response to different amount of 7-tricosene or to the solvent only (hexane). (C) Ejaculate ejection latency female *CS* in response to different amount of heptanal or to the solvent only (hexane). Numbers in between brackets indicate sample sizes. mean ± S.E.M are indicated in the graphs. Scatter-plot labelled with the same letter are not statistically different. *n.s*.: non-significant; *:p<0.05; **:p<0.01; ***:p<0.001.For full statistical details see Supplementary Table S1.

